# An Excelled Trifluorinated Probe for GPCR Conformational Quantification

**DOI:** 10.64898/2026.09.18.752616

**Authors:** Xudong Wang, Yulong Cao, Wenjie Zhao, Wenkai Sun, Trang Ma, Yongbo Zhang, Libin Ye

**Author notes:** **Corresponding authors Libin Ye, Associate Professor,** Department of Molecular Biosciences, University of South Florida, 4202 E Fowler Ave, Tampa, FL, USA 33620; H. Lee Moffitt Cancer Center & Research Institute, 12902 USF Magnolia Drive, Tampa, FL, USA 33612.

## Abstract

G protein-coupled receptors (GPCRs) represent the largest membrane protein family in the human body and the major targets of FDA-approved medications. Unlike some other physiological proteins, GPCRs are highly plastic and often exist as collections of conformational ensembles in the environment to exert their function. Therefore, understanding how a GPCR adopts different conformational ensembles in response to ligands and signaling partners is essential for drug development. ^19^F-NMR has emerged as one of the most promising tools in this regard; however, a well-established ^19^F-labeling system is still lacking, including an effective ^19^F probe that meets both environmental susceptibility and NMR sensitivity requirements, even though progress has been made dramatically in recent years. Thus, this manuscript reports a novel conformational probe (4-(bromomethyl)- ethoxy-2-(trifluoromethyl)benzene, BTMB) that offers paramount benefits for future GPCR conformational dynamics studies. To rationalize the mechanism behind BTMB’s superiority for future probe development, we performed MD simulations, referenced to the current state-of-the-art probe, BTFMA. Our study suggests that the spatial proximity of aromatic residues significantly influences the chemical shift dispersion and linewidths of related ^19^F-probe-profiled conformational states, as well as the rigidity and compactness of -CF_3_ in the target protein.

## Introduction

High-resolution structures have defined inactive, agonist-bound, and G-protein-coupled states of the GPCR, including the adenosine A_2A_ receptor (A_2A_R), in which activation of the GPCR leads to outward displacement of transmembrane domain 6 (TM6), opening the intracellular G protein binding pocket. A set of biophysical tools such as NMR, DEER, and FRET has extended these static structural snapshots by showing continuous conformational re-equilibration among a set of states in response to the binding of inverse, partial, and full agonists, as well as G proteins^1–5^. However, even when bound to both a high-affinity ligand and the cognate downstream heterotrimeric G proteins, the receptor doesn’t completely present as a single conformational state, but as a conformational ensemble with a redistributed subpopulation of each state^6–8^. This highlights the importance of developing tools to visualize the receptor’s continuous conformational landscape and to study how it responds to ligands and signaling partners.

As mentioned earlier, despite many tools being underdeveloped for probing conformational states, ^19^F NMR stands out as one of the most powerful options because fluorine offers high sensitivity to micro-electrostatic environmental changes and has minimal biological background ^9–15^. So far, using ^19^F-NMR via a site-specific fluorine attachment, several GPCRs have been resolved in the inactive, intermediate, and active states, as well as their transitions among different states^2–5,9–13,16,17^. One example is the adenosine 2A receptor (A_2A_R) using BTFMA labeling at V229C, a mobile site on transmembrane domain 6. Using this system as a reference, we report a novel trifluorinated tag, BTMB, that exhibits a much more sensitive response to receptor motion, triggered by ligands or G proteins^2,3^.

The static electronic environment usually consists of electrostatics, van der Waals packing, solvent exposure, probe rotamers, and aromatic ring currents surrounding the ^19^F probe ^18–22^. To assess BTMB’s sensitivity to the static electronic environment, we performed a series of 100 ns molecular dynamics (MD) simulations to probe its aromatic contact geometry, as two aromatic residues (F201^5.62^ and H230^6.32^) are nearby^23^. We expect to connect the ^19^F-NMR spectral feature to a state-dependent microenvironment surrounding the ^19^F reporter, which should guide future probe discovery; probe discovery is a continued, ongoing, never-ending task, and we expect to develop an even better probe to study protein conformational dynamics in the future, guided by the method and progress presented here.

## Results

### Compound BTMB showed an explicit response to the microenvironment

Using a methanol/water gradient to vary microenvironment polarity^24^, we examined 38 aryl-CF_3_ compounds (Supplemental Table 1) for their susceptibility by acquiring 1D ^19^F NMR. We calculated the chemical shift deviation Δδ to assess their susceptibility to solvent polarity changes. As shown in Fig. 1, BTMB exhibited the largest chemical shift amplitude among the compounds, including the state-of-the-art probe BTFMA, with Δδ of 1.71 ppm (BTMB) vs 0.95 ppm (BTFMA). Of note, monofluoro-compounds usually exhibit greater microenvironmental sensitivity due to a smaller chemical shift anisotropy effect^25,26^. However, we don’t discuss this further in this manuscript because we expect trifluoride to offset the low-yield expression of GPCRs, ensuring that the developed ^19^F probe has broad applicability in membrane protein research. Because the atomic geometries of BTMB and BTFMA differ slightly, their substitution patterns differ when conjugated to V229C, a labeling site on transmembrane domain 6. BTMB forms a relatively compact Cys–S–CH₂–Ar benzyl-thioether linkage, whereas BTFMA forms a longer and more flexible Cys–S–CH₂–C(O)–NH–Ar linkage. This structural difference could contribute to the linewidth distinction observed for these two probes when conjugated to the receptors. We will discuss this further in the follow-up sections.

**Figure 1.**
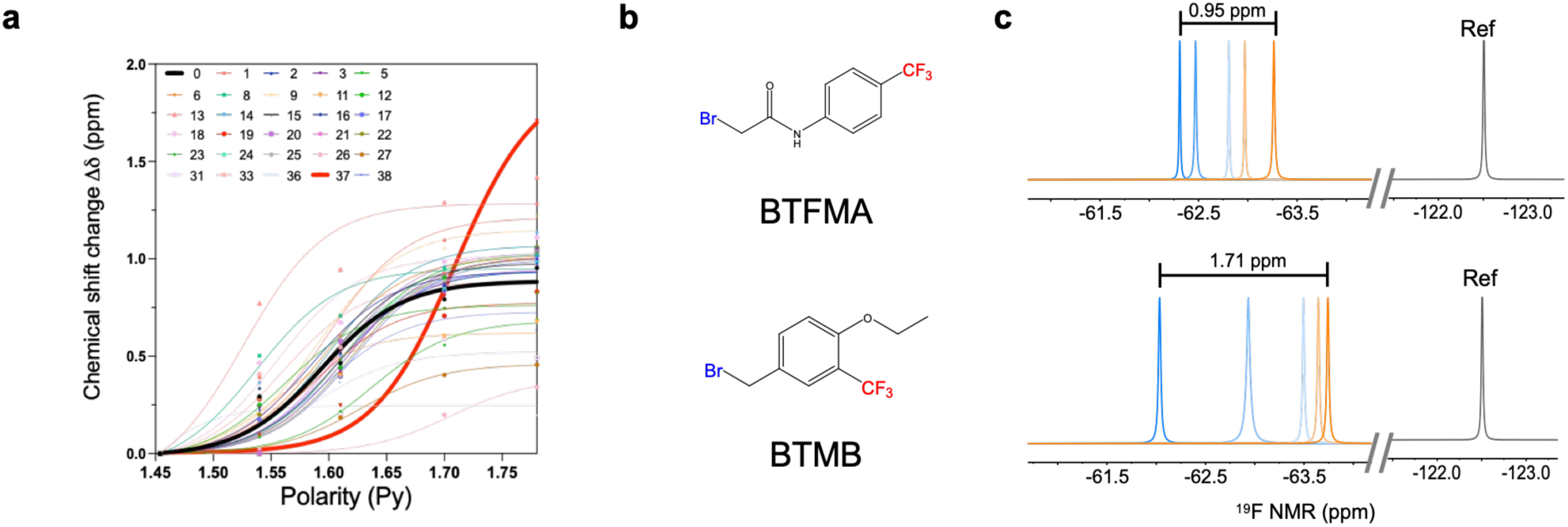
A comparison of chemical shift dispersion in response to solvent polarity changes. a, Changes in 19F chemical shift (Δδ) of various -CF3 tags as a function of solvent polarity. Note: that all changes are referenced to the most hydrophobic solution investigated (i.e. MeOH/H_2_O = 4.0 or Py = 1.454). Observed changes in chemical shift were fit to the formula Δδ = *A*/(1 + exp(−(Py − *x _0_*)/*w*)), where *A*, *x _0_*, and *w* represent fitting parameters for each probe. b. Chemical structures of the reference BTFMA and compound BTMB. c. Representative 19F NMR spectra of BTFMA (top panel) and BTMB (bottom panel), and their comparison in chemical shift dispersion as a function of solvent polarity.

**Supplemental Table 1:**
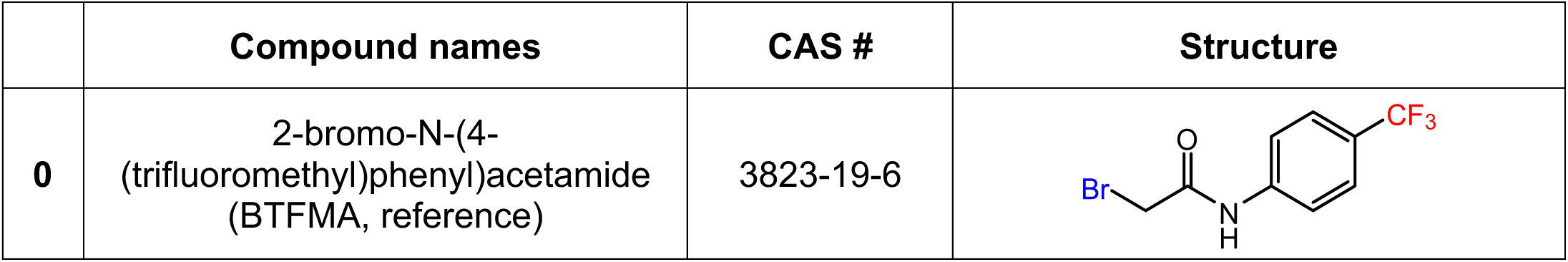
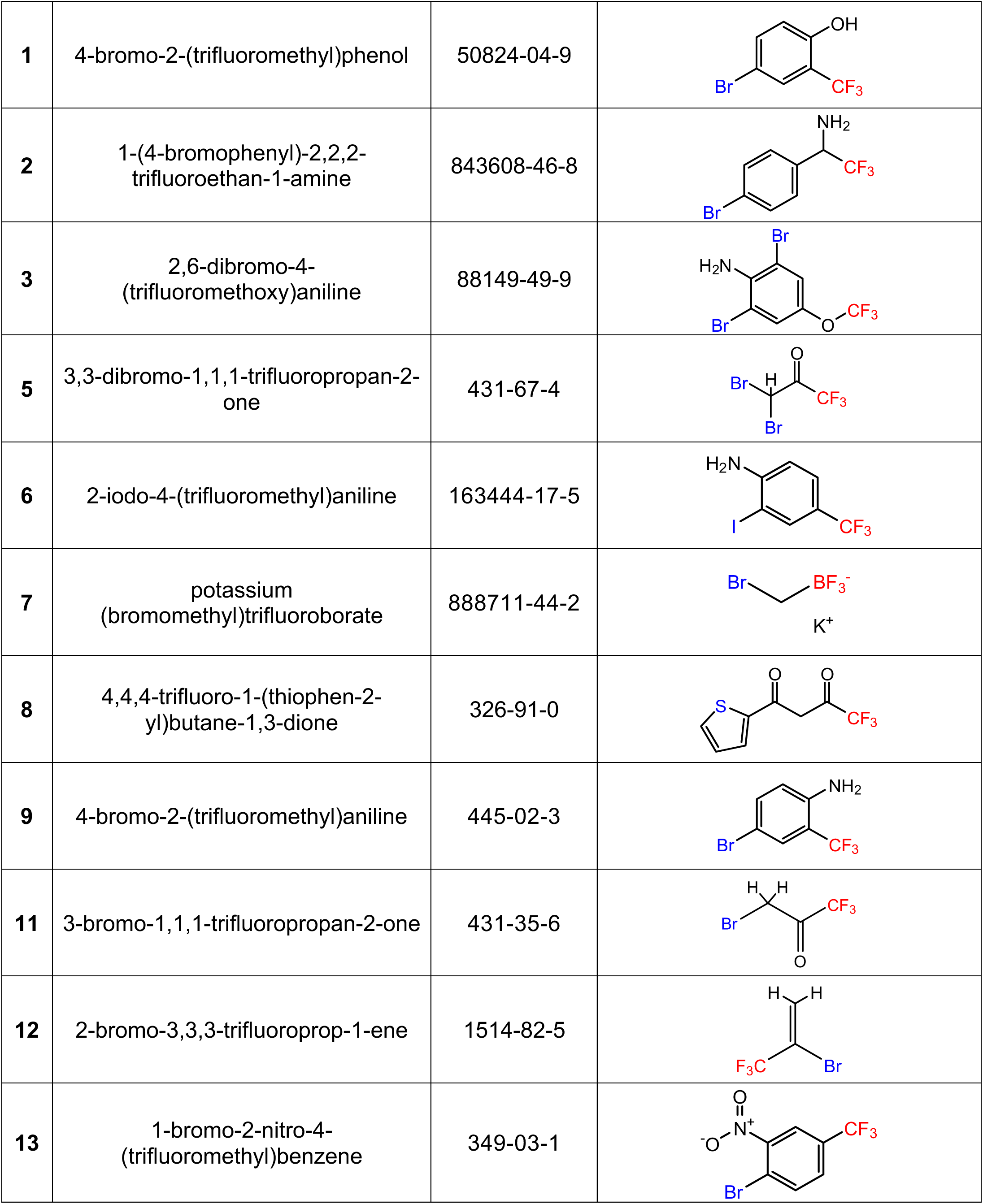
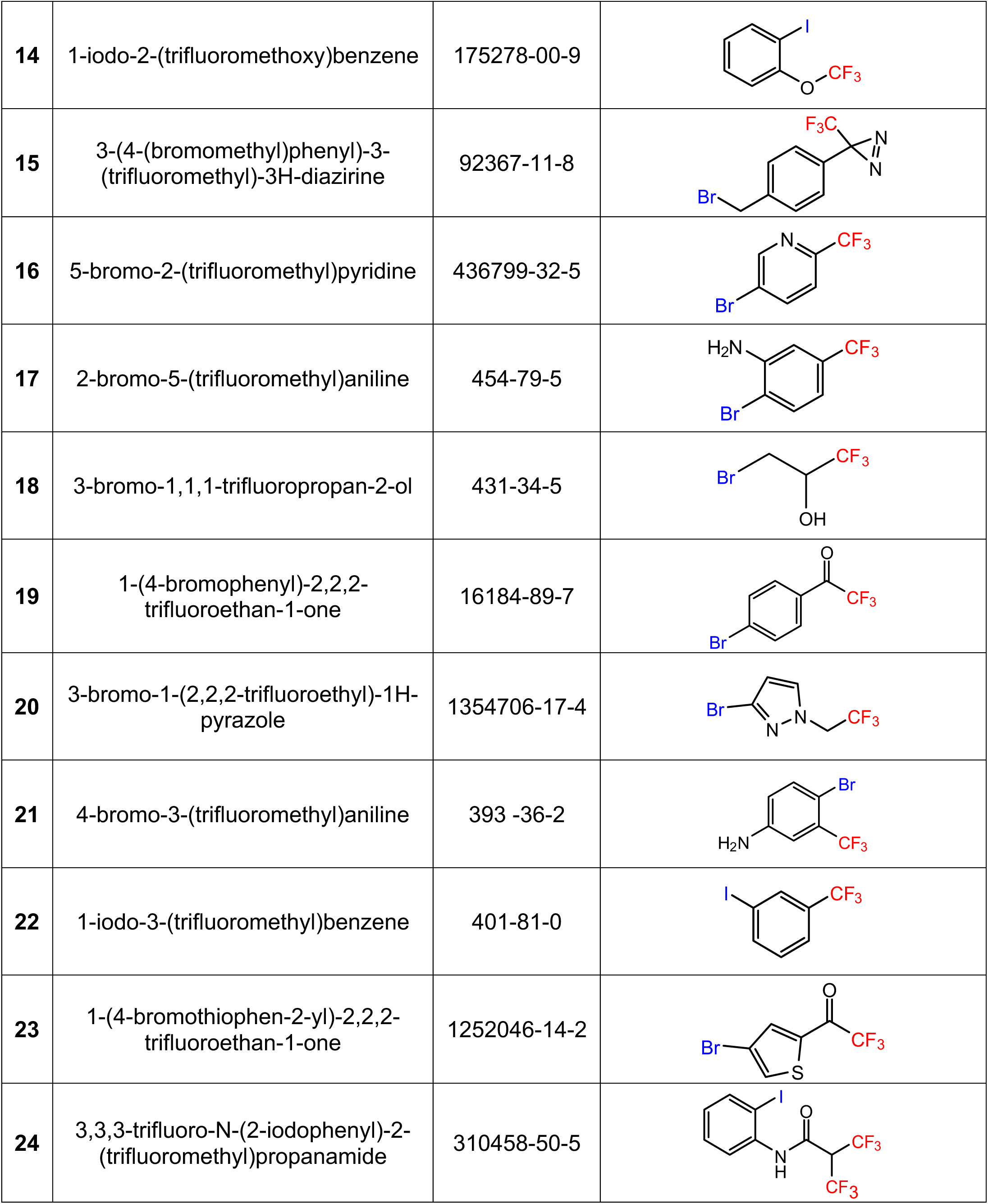
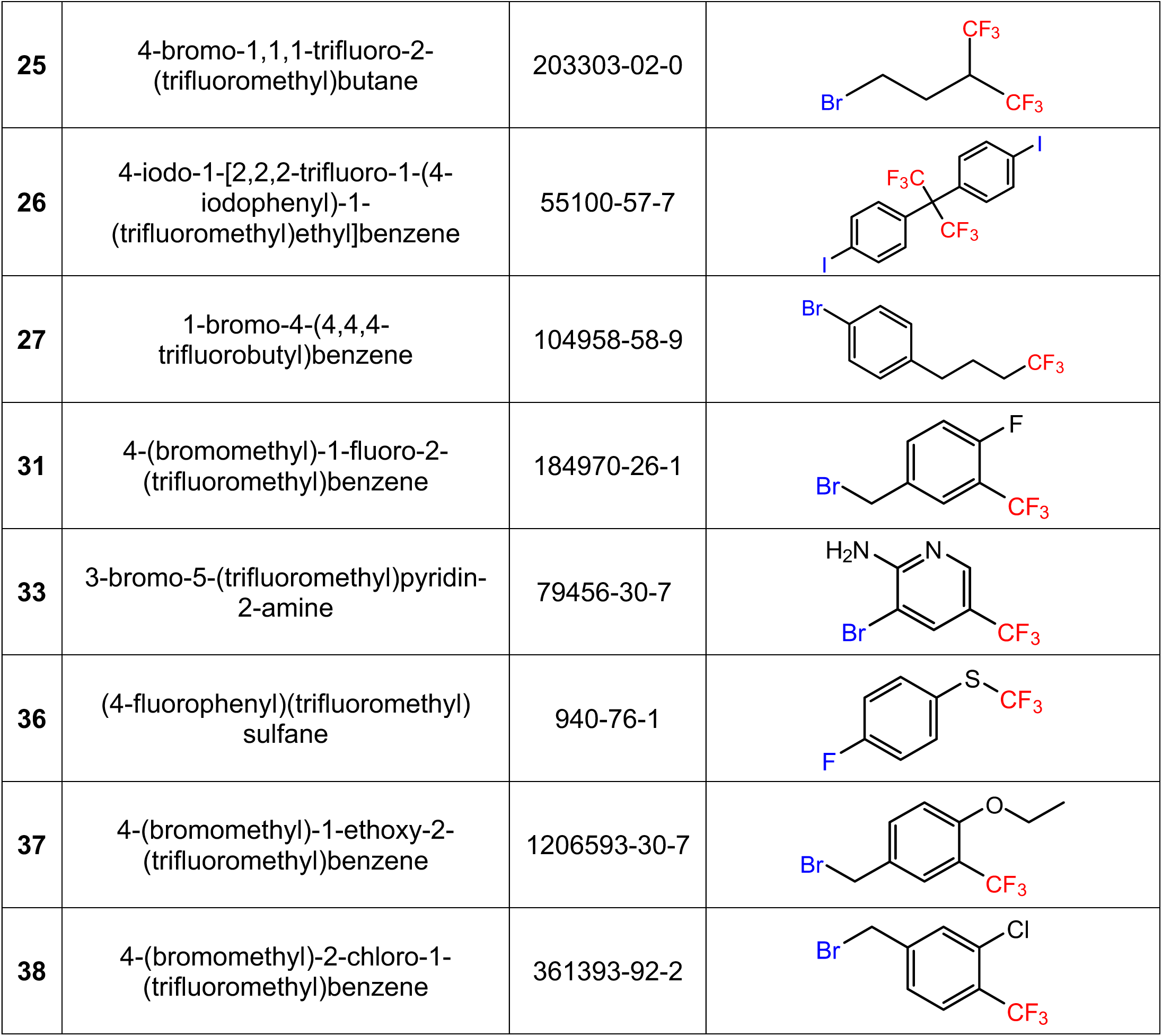
Trifluorinated compounds used in this study.

|  | Compound names | CAS # | Structure |
| --- | --- | --- | --- |
| **0** | 2-bromo-N-(4-(trifluoromethyl)phenyl)acetamide (BTFMA, reference) | 3823-19-6 |  |
| 1 | 4-bromo-2-(trifluoromethyl)phenol | 50824-04-9 |  |
| 2 | 1-(4-bromophenyl)-2,2,2-trifluoroethan-1-amine | 843608-46-8 |  |
| 3 | 2,6-dibromo-4-(trifluoromethoxy)aniline | 88149-49-9 |  |
| 5 | 3,3-dibromo-1,1,1-trifluoropropan-2-one | 431-67-4 |  |
| 6 | 2-iodo-4-(trifluoromethyl)aniline | 163444-17-5 |  |
| 7 | potassium (bromomethyl)trifluoroborate | 888711-44-2 |  |
| 8 | 4,4,4-trifluoro-1-(thiophen-2-yl)butane-1,3-dione | 326-91-0 |  |
| 9 | 4-bromo-2-(trifluoromethyl)aniline | 445-02-3 |  |
| 11 | 3-bromo-1,1,1-trifluoropropan-2-one | 431-35-6 |  |
| 12 | 2-bromo-3,3,3-trifluoroprop-1-ene | 1514-82-5 |  |
| 13 | 1-bromo-2-nitro-4-(trifluoromethyl)benzene | 349-03-1 |  |
| 14 | 1-iodo-2-(trifluoromethoxy)benzene | 175278-00-9 |  |
| 15 | 3-(4-(bromomethyl)phenyl)-3-(trifluoromethyl)-3H-diazirine | 92367-11-8 |  |
| 16 | 5-bromo-2-(trifluoromethyl)pyridine | 436799-32-5 |  |
| 17 | 2-bromo-5-(trifluoromethyl)aniline | 454-79-5 |  |
| 18 | 3-bromo-1,1,1-trifluoropropan-2-ol | 431-34-5 |  |
| 19 | 1-(4-bromophenyl)-2,2,2-trifluoroethan-1-one | 16184-89-7 |  |
| 20 | 3-bromo-1-(2,2,2-trifluoroethyl)-1H-pyrazole | 1354706-17-4 |  |
| 21 | 4-bromo-3-(trifluoromethyl)aniline | 393 -36-2 |  |
| 22 | 1-iodo-3-(trifluoromethyl)benzene | 401-81-0 |  |
| 23 | 1-(4-bromothiophen-2-yl)-2,2,2-trifluoroethan-1-one | 1252046-14-2 |  |
| 24 | 3,3,3-trifluoro-N-(2-iodophenyl)-2-(trifluoromethyl)propanamide | 310458-50-5 |  |
| 25 | 4-bromo-1,1,1-trifluoro-2-(trifluoromethyl)butane | 203303-02-0 |  |
| 26 | 4-iodo-1-[2,2,2-trifluoro-1-(4-iodophenyl)-1-(trifluoromethyl)ethyl]benzene | 55100-57-7 |  |
| 27 | 1-bromo-4-(4,4,4-trifluorobutyl)benzene | 104958-58-9 |  |
| 31 | 4-(bromomethyl)-1-fluoro-2-(trifluoromethyl)benzene | 184970-26-1 |  |
| 33 | 3-bromo-5-(trifluoromethyl)pyridin-2-amine | 79456-30-7 |  |
| 36 | (4-fluorophenyl)(trifluoromethyl) sulfane | 940-76-1 |  |
| 37 | 4-(bromomethyl)-1-ethoxy-2-(trifluoromethyl)benzene | 1206593-30-7 |  |
| 38 | 4-(bromomethyl)-2-chloro-1-(trifluoromethyl)benzene | 361393-92-2 |  |

### BTMB labeling and structure-functional perturbation assessment

BTMB’s distinctive response to solvent polarity prompted us to assess its suitability for receptor applications. Using a well-established GPCR system in the lab—the adenosine 2A receptor (A_2A_R) — and a judiciously selected labeling site that clearly reflects the receptor’s conformational ensemble through BTFMA attachment in our early studies, we aim to examine whether the new tag outperforms BTFMA^2,5^. We conducted a series of experiments to assess labeling efficiency, the effects of labeling on structural and functional perturbations, and the functional outcomes of A_2A_R_V229C^6.31^_BTMB in the A_2A_R transmembrane domain VI (TM6), a site that responds elegantly to conformational changes during receptor activation^2,5,17^.

As illustrated in Fig.2A, BTMB is conjugated to the V229C^6.31^ residue via the thiol group from the cysteine residue. We confirmed successful conjugation by MS (Supplemental Figure 1). Unlike BTFMA, which contains an α-bromoacetamide group (Ar–NH–C(O)–CH₂–Br), BTMB conjugates to cysteine through a benzylic bromide group (Ar–CH₂–Br). Although both α-bromoacetamides and benzylic bromides undergo nucleophilic substitution with thiolates, α-bromoacetamides are well-established cysteine-reactive electrophiles that typically undergo rapid S_N2 alkylation^27,28^, whereas benzylic bromides represent a chemically distinct class of electrophiles whose reactivity is governed by the benzylic environment and aromatic substitution^29,30^. Because GPCRs are time-sensitive receptors, we evaluated BTMB labeling efficiency via Ellman’s assay ^31^ (Fig. 2c), in comparison with BTFMA, to determine and optimize its application for GPCR labeling. As mentioned above, the data indeed indicated that the labeling efficiency of BTMB is much lower than BTFMA at 4 °C at the concentration of 50 μM, but a complete labeling can be achieved in 20 h with 200 μM BTMB (Fig.2c). This indicates it’s suitable for GPCR use without impairing function. Our further experiments also showed that the BTMB-labeled receptor was fully comparable to the WT construct and the BTFMA-labeled receptor in regulating heterotrimeric G proteins to hydrolyze GTP (Fig. 2d). These results strongly suggest that BTMB is a promising ^19^F reporter for time-sensitive proteins, suitable for GPCR labeling without functional disruption, enabling broad application.

**Figure 2.**
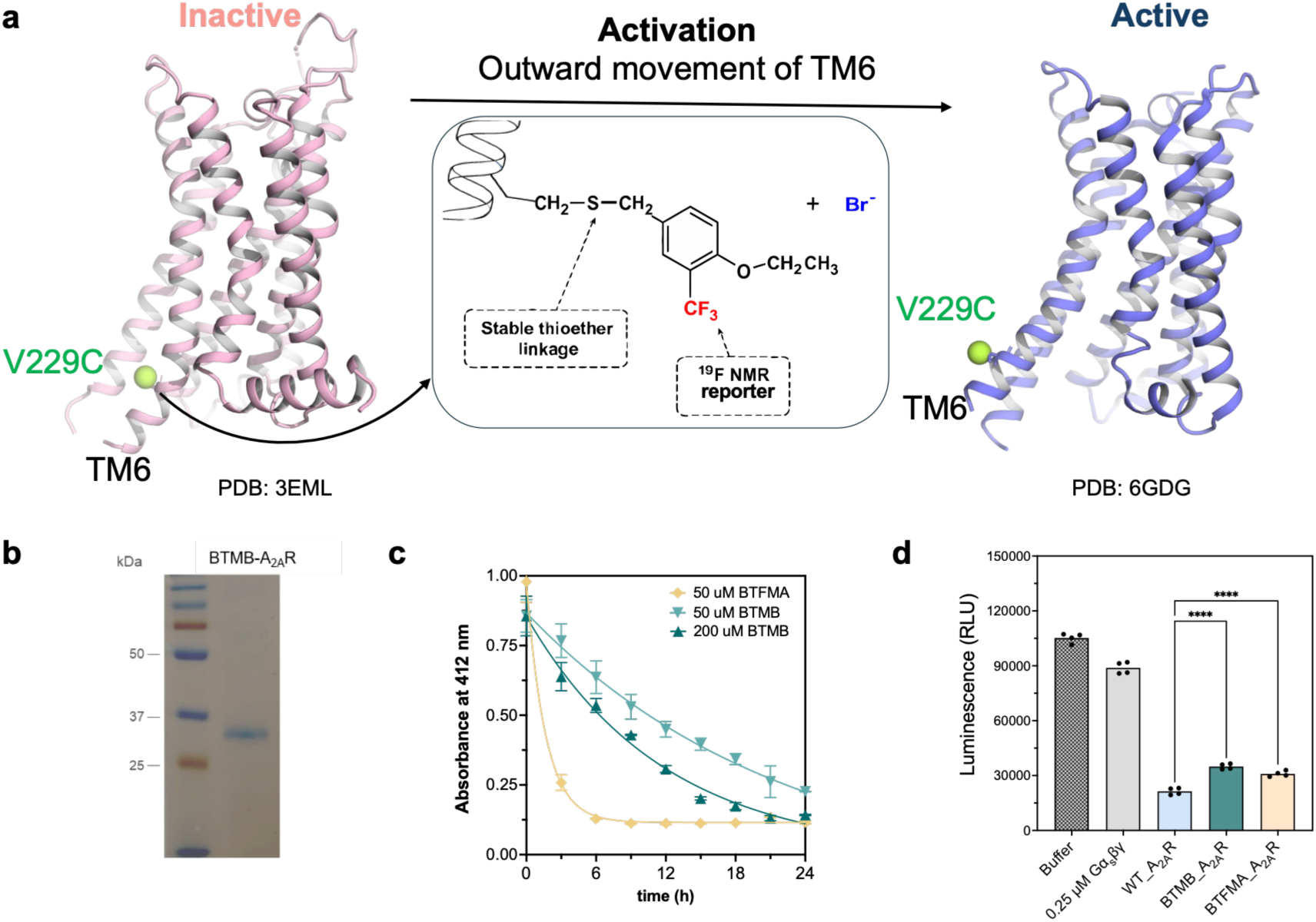
Biochemical and functional characterization of BTMB labeling and comparison to the state-of-the-art probe BTFMA. **a**. Topology of BTMB conjugation on V229C in the A_2A_R receptor on the TM6 domain, presented in the inactive (PDB ID: 3EML) and active (PDB ID: 6GDG) states. **b**. SDS-PAGE of BTMB-conjugated A_2A_R receptor. **c**. Labeling efficiency assessments of BTMB and BTFMA conjugations to the receptor. **d**. GTP hydrolysis capacity of BTMB- and BTFMA-conjugated A_2A_R in comparison to the WT.

**Supplemental Figure 1.**
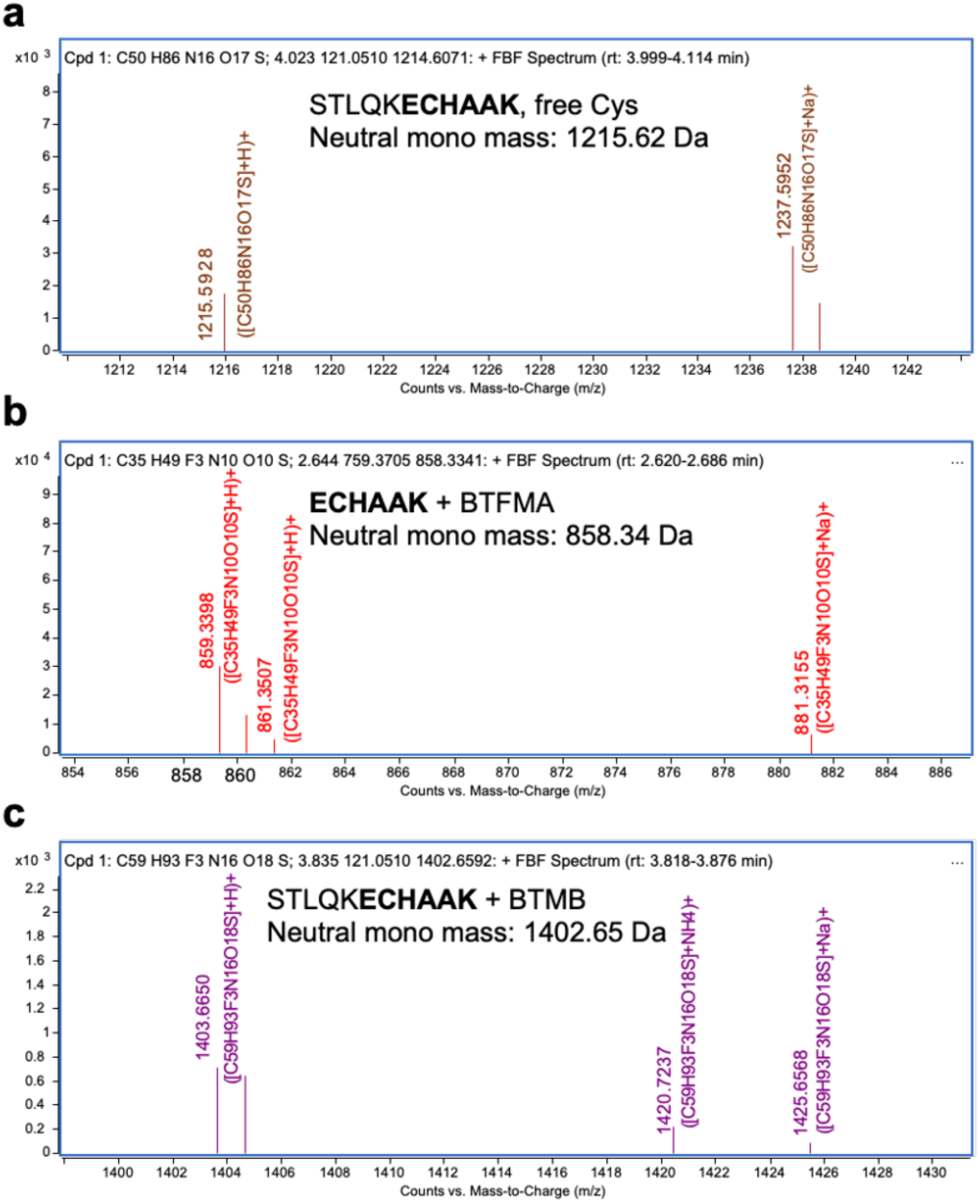
LC–MS identification of unmodified and probe-modified A2AR-V229C peptides. **a,** Representative positive-ion LC–TOF MS spectrum of the unmodified A_2A_R-V229C peptide species detected after tryptic digestion, corresponding to C₅₀H₈₆N₁₆O₁₇S at a retention time of 3.999–4.114 min, with representative signals at *m/z* 1215.5928 and 1237.5952. **b**, Representative spectrum of the BTFMA-modified peptide species, corresponding to C₃₅H₄₉F₃N₁₀O₁₀S at 2.620–2.686 min, with representative signals at *m/z* 859.3398 and 881.3155. **c**, Representative spectrum of the BTMB-modified peptide species, corresponding to C₅₉H₉₃F₃N₁₆O₁₈S at 3.818–3.876 min, with representative signals at *m/z* 1403.6660 and 1425.6568. Major peaks include protonated and sodium-adducted ions.

### BTMB possesses superiority over BTFMA in profiling GPCR dynamic conformations

Given the success of BTMB labeling and its ability to maintain function, we aim to evaluate how well BTMB profiles GPCR conformational dynamics, using BTMB-labeled A_2A_R_V229C as the evaluation system, in reference to that of BTFMA. We acquired ^19^F NMR spectra as a function of ligand and performed spectral deconvolution to assess the A_2A_R conformational landscape. As shown in Fig.3, we were able to create a conformational matrix to delineate the A_2A_R into three major conformational ensemble clusters, corresponding to inactive, partially activated, and fully activated conformations in response to various ligands. Each ensemble can be further delineated into 2-3 substates featured with different colors, from red (P1 and P2), yellow (P3, P4, and P5), to blue (P6, P7, and P8), guided by T2 measurements (Supplemental Figure 2). In contrast, in our previous studies on BTFMA-labeled A_2A_R, we couldn’t spectroscopically distinguish these substates^2,17^. Together, our results demonstrate BTMB’s sensitivity in dissecting and quantifying the receptor’s conformational states, with superior performance over BTFMA, including larger chemical-shift dispersion (819Hz vs 774 Hz) (Supplemental Figure 3a) and narrower linewidths for different components (Supplemental Figure 3b).

**Supplemental Figure 2.**
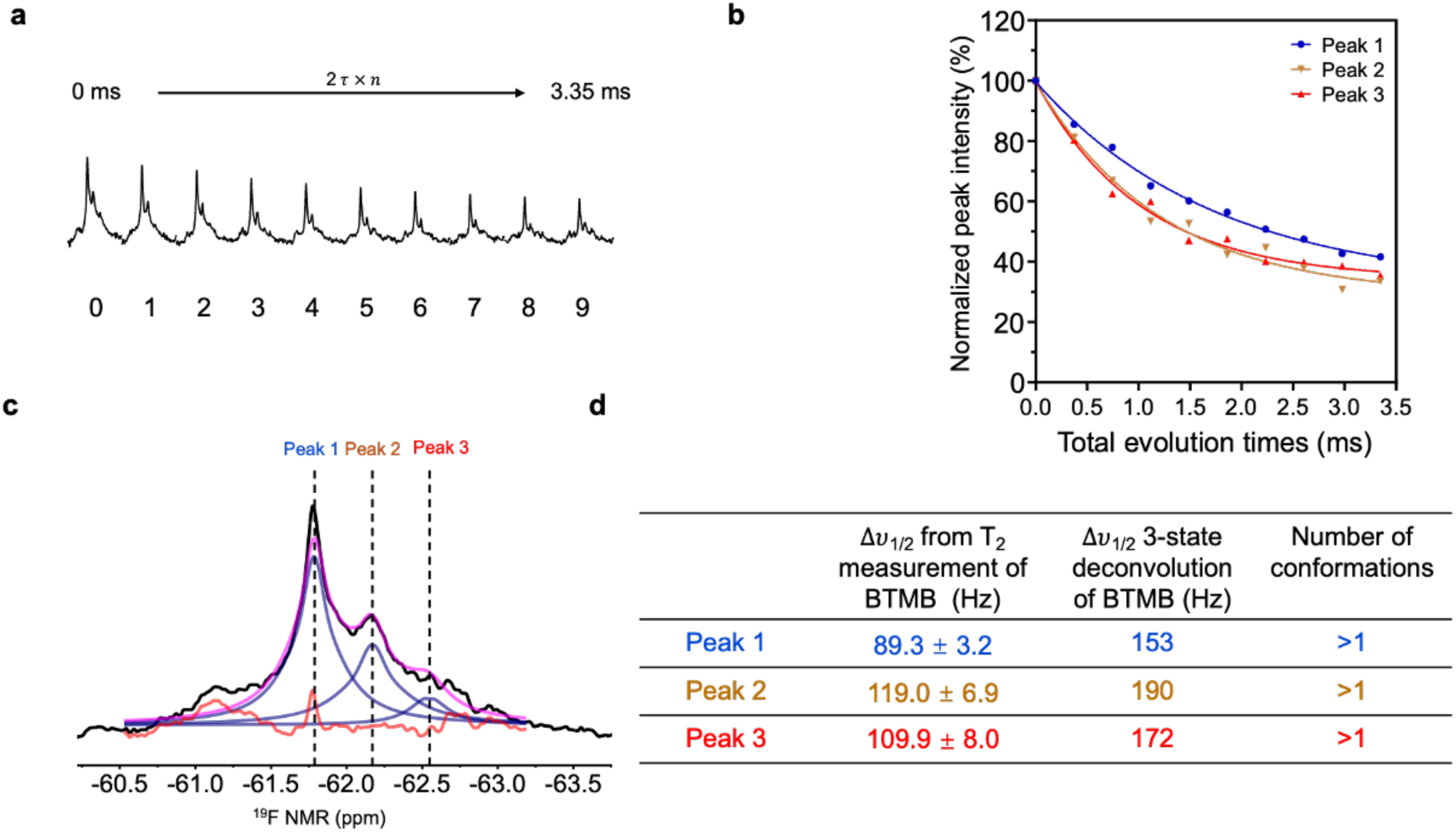
^19^F NMR CPMG relaxation measurements of the BTMB-labeled A_2A_R to determine the linewidths of each conformational component. **a,** T2 relaxation experiments for linewidth measurements of different resonances. **b,** T2 relaxation fitting and the corresponding linewidth from the fitting shown in **(d)**. **c,** 2-state based spectral deconvolution for the major resonance that consists of Peak 2 and Peak3. **D,** The linewidths of each deconvoluted resonance.

**Figure 3.**
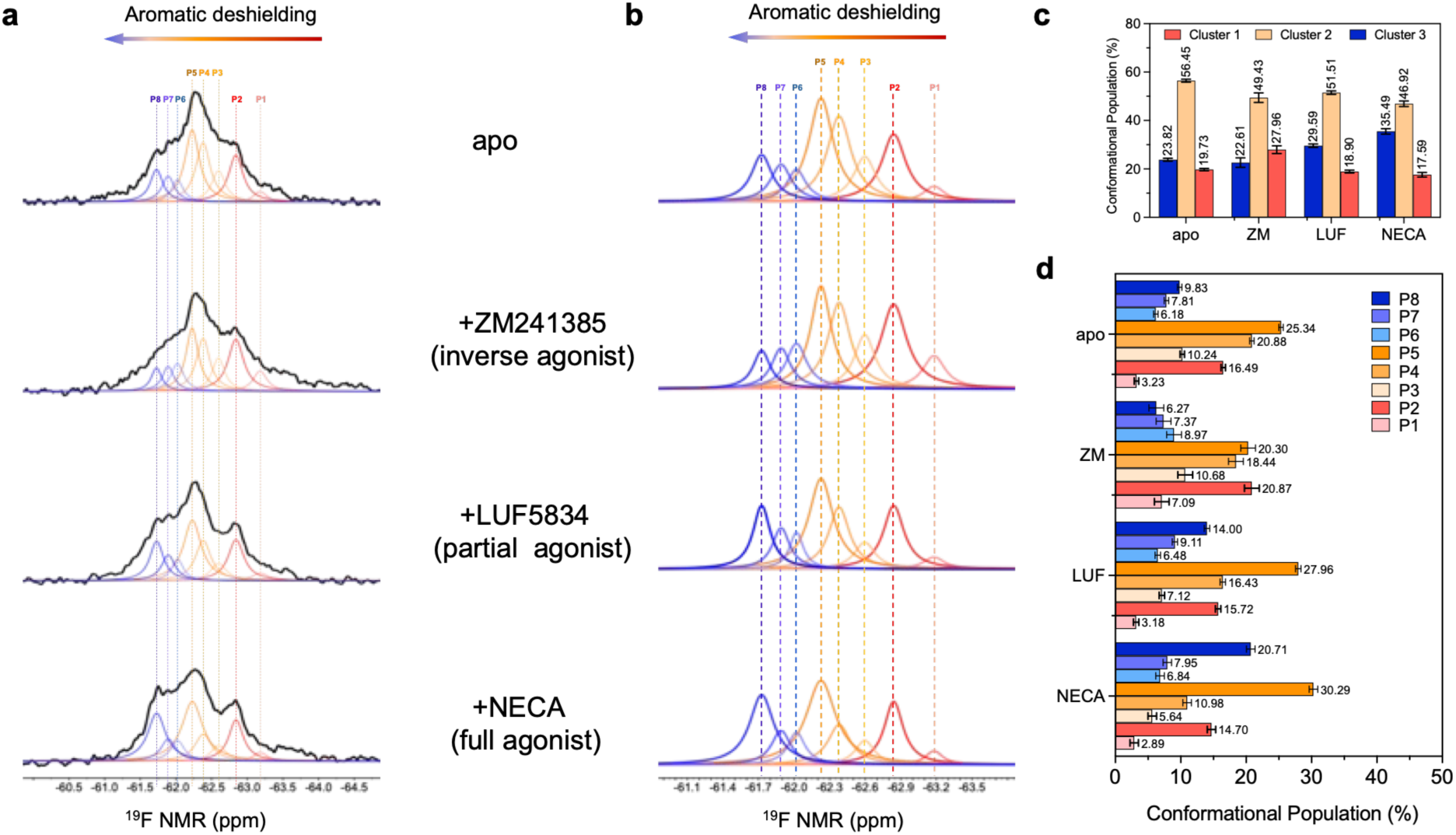
Defined conformational landscapes of the A_2A_R analyzed with BTMB labeling. **a.** Conformational landscape of the A_2A_R as a function of ligand binding, analyzed using BTMB labeling at V229C in the TM6 domain. **b.** Deconvoluted BTMB-labeled A_2A_R as a function of ligand binding. **c**. Histograms of the 3 clustered conformational ensembles in response to different ligand binding. **d.** Histograms of the subpopulations of each delineated conformational state in response to different ligand binding.

**Supplemental Figure 3.**
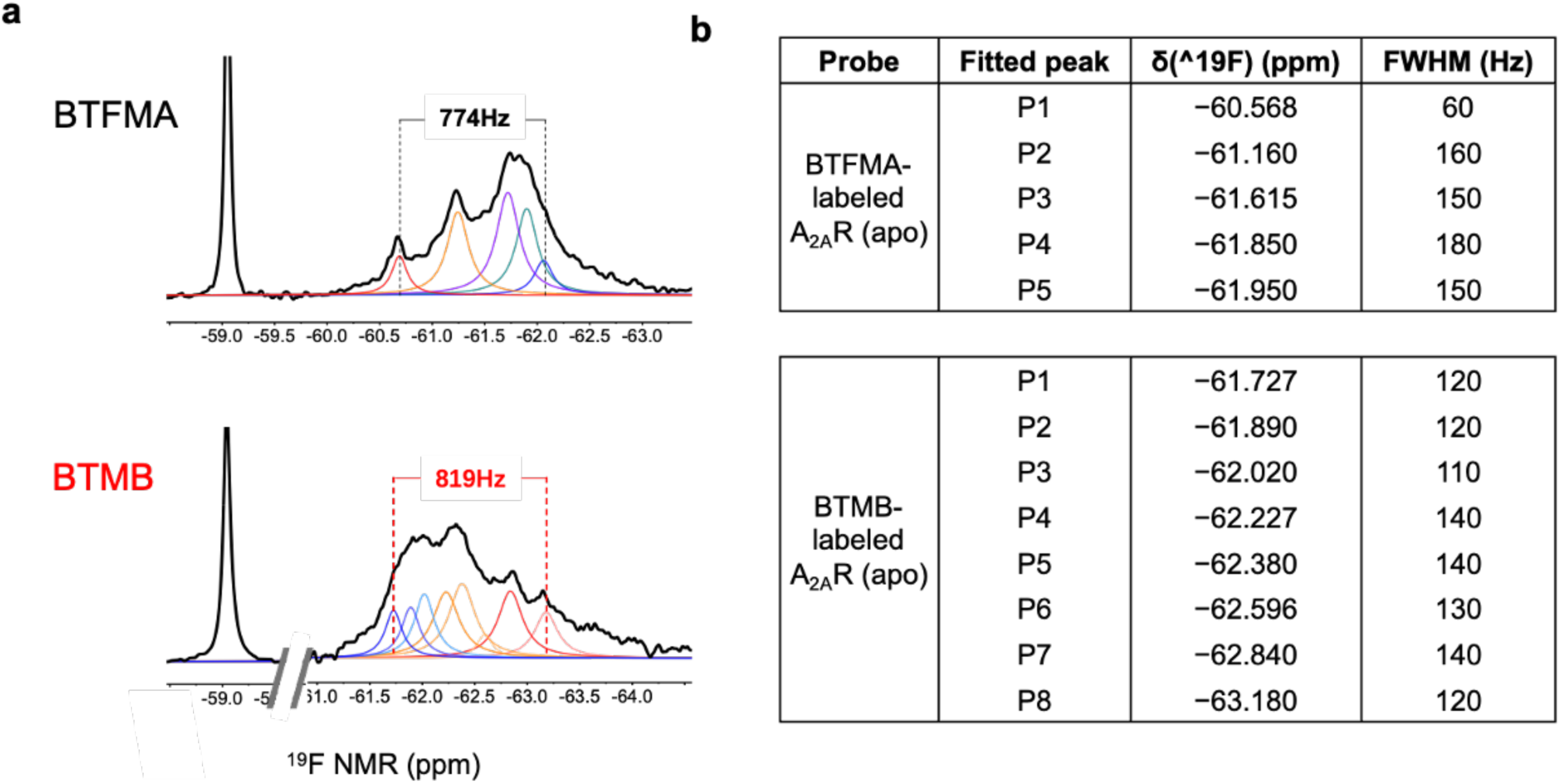
Comparison of fitted ^19^F NMR resonances and chemical-shift dispersion for BTFMA- and BTMB-labeled A_2A_R. **a,** Comparison of the overall chemical-shift dispersion of BTFMA- and BTMB-labeled A2AR, using the most distant fitted resonances as boundaries. The corresponding dispersions were approximately 774 Hz for BTFMA and 819 Hz for BTMB. Individual fitted components are shown in different colors. **b,** Summary of the fitted resonance positions and corresponding full widths at half-maximum (FWHM) obtained from the **apo** spectra of BTFMA- and BTMB-labeled A_2A_R.

In addition to performing experiments with BTMB-labeled A_2A_R as a function of ligand, we also tested whether adding G proteins further drives conformational re-equilibration of BTMB-labeled A_2A_R toward the fully activated state. As illustrated in Fig. 4, the presence of heterotrimeric G protein prominently shifted the A_2A_R conformational ensemble toward the P8 state, at the expense of inactive and partially activated states. This also indicates that the BTMB-labeled GPCR, A_2A_R, responds not only to the ligand but also to downstream signaling partners such as G protein engagement, further demonstrating that the labeling retains receptor functionality with minimal structure-function perturbation. At this point, we can conclude that compound BTMB demonstrates superior responsiveness to microenvironmental changes as a next-generation ^19^F probe for studying conformational dynamics of proteins like GPCRs. Although its labeling efficiency is slightly less than BTFMA’s, this can be offset by using a multiple-labeling strategy at higher concentrations, as shown in (Fig.2c).

**Figure 4.**
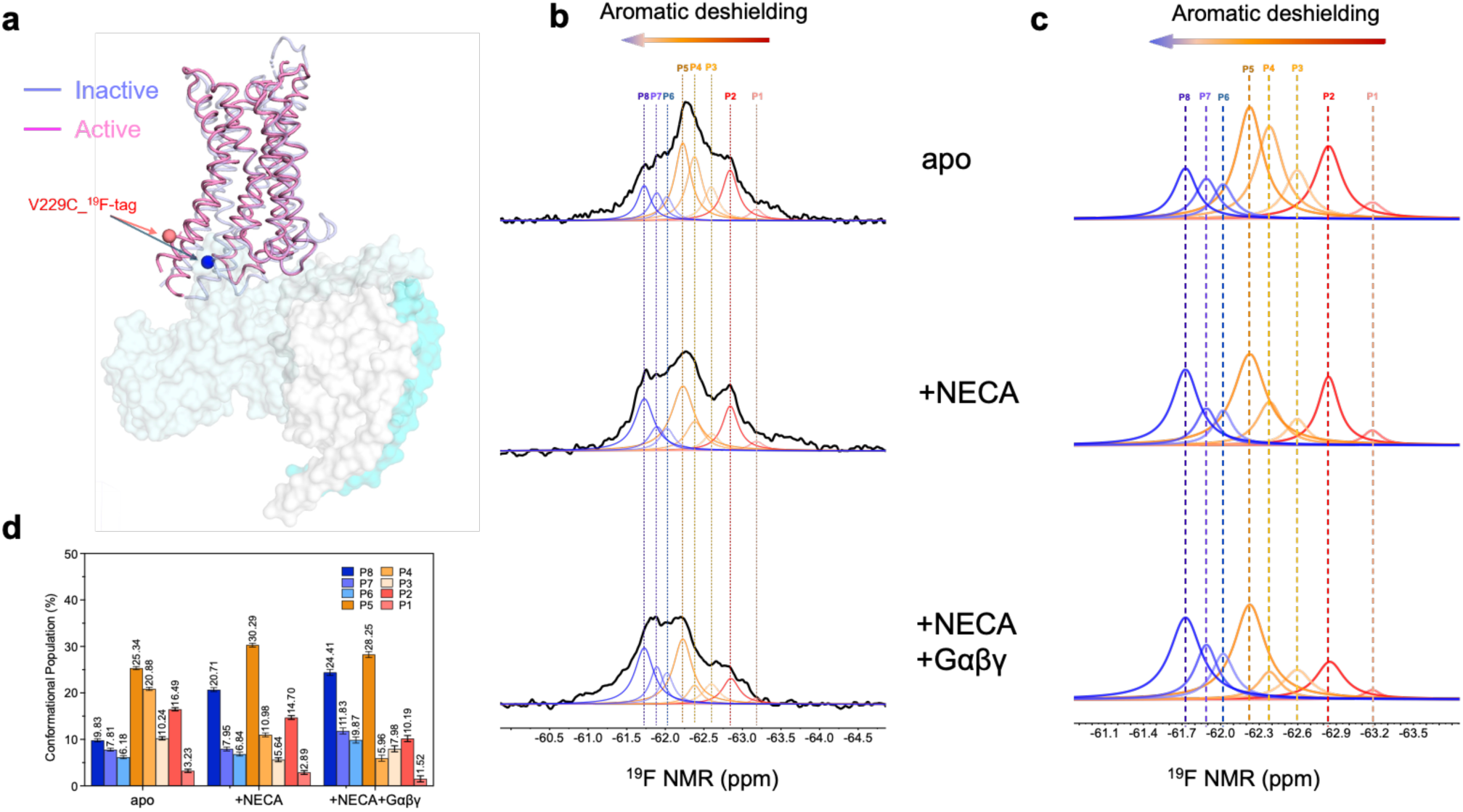
Population analysis of BTMB -resolved A_2A_R conformational ensembles. **a,** Structural comparison of inactive (blue) and active (magenta) A_2A_R conformations showing the V229C_ ^19^F reporter site at the intracellular end of TM6. **b,** Representative deconvoluted ^19^F NMR spectra of BTMB - labelled A_2A_R under the indicated conditions, shown at two horizontal scales to display the overall spectral distribution (middle) and (**c)** the overlapping fitted components (right). Eight resonances are color-coded consistently across conditions. **d,** Relative populations of the fitted P1–P8 resonances under the indicated conditions. Peak areas were normalized to the total fitted ^19^F signal for each spectrum; error bars represent the signal-to-noise ratio of the corresponding resonances.

### Potential mechanism of the superiority of BTMB over BTFMA

To rationalize the molecular mechanism behind BTMB-A_2A_R superiority over BTFMA-A_2A_R, we simulated both probes in the inactive and active states of the A_2A_R receptor using the ZM- and NECA-bound receptor structures: PDB 3EML (inactive) and 6GDG (fully activated), respectively^32,33^. Each ligand-bound system was run for 100 ns, and 2,000 evenly spaced frames were analyzed for aromatic contacts surrounding the CF_3_ group.

Two prominent neighboring aromatic residues alternate with the receptor state (Figs. 5a and 5b). In the ZM-bound inactive-start trajectory, F201^5.62^ remained persistently proximal to BTMB (4.45 ± 0.40 Å; 99.6% of frames within 6 Å), whereas in the NECA-bound active-start trajectory it was substantially farther away (12.70 ± 0.45 Å). Conversely, H230^6.32^ showed the opposite pattern: it remained distant in the ZM-bound trajectory (11.72 ± 0.92 Å) but remained close in the NECA-bound active-start trajectory (5.24 ± 0.39 Å; 96.5% of frames within 6 Å). In the case of BTFMA-labeled A_2A_R, the effects of these two aromatic residues on the -CF_3_ group seemed relatively mild because BTFMA has a longer linker to the receptor, leading to the CF_3_ group being located slightly farther from these two aromatic residues (Figs. 5c and 5d) and thus reducing its influence from the aromatic residues F201^5.62^ and H230^6.32^.

**Figure 5.**
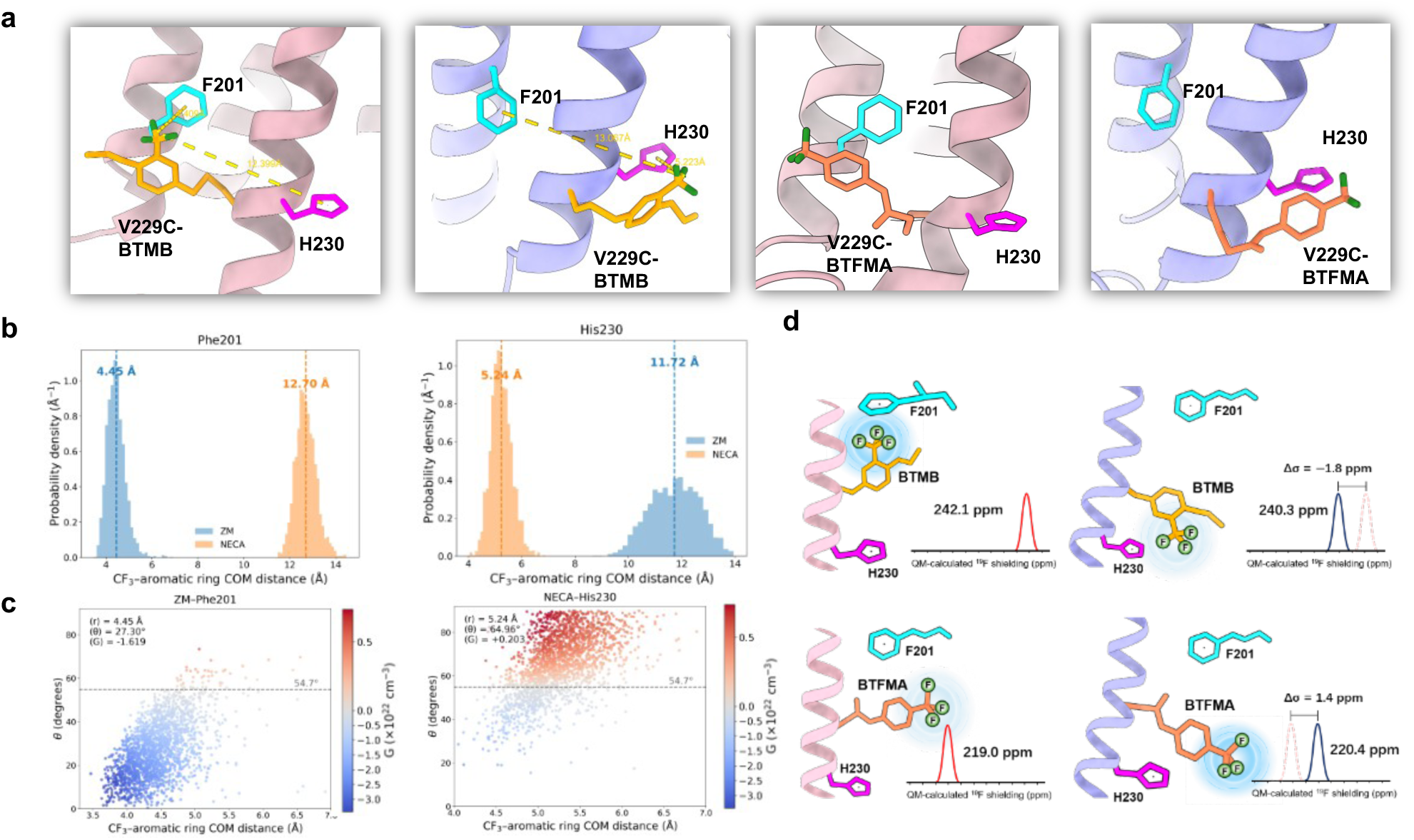
Structural, geometric, and quantum-chemical analysis of the surrounding environment of the ^19^F_A_2A_R_V229C. **a,** Representative poses of BTMB - and BTFMA-labelled A_2A_R_V229C^6.31^ in the ZM- and NECA-bound receptor. The ZM- and NECA-bound receptor conformations are shown in pink and blue, respectively; the ^19^F probes are shown in orange, F201^5.62^ in cyan, and H230^6.32^ in magenta. **b,** Probability-density distributions of the CF_3_-to-aromatic-ring centre-of-mass distances for F201^5.62^ and H230^6.32^ in the BTMB simulations. ZM and NECA trajectories are shown in blue and orange, respectively. **c,** Joint distributions of the CF_3_-to-aromatic-ring centre-of-mass distance, r, and the orientation angle, theta, for the indicated aromatic residues. Theta is defined by the aromatic-ring normal and the vector connecting the aromatic-ring centre to the CF_3_ group. Each point represents a simulation frame and is coloured according to the geometric ring-current factor G. **d,** Schematic representation of the local CF_3_ environments of BTMB and BTFMA in the ZM- and NECA-bound models, together with QM-calculated ^19^F isotropic shielding.

Ring-current effects of aromatic residues depend strongly on both orientation and distance; we therefore calculated G = (1 - 3cos^2^θ)/r^3^, where θ is the angle between the aromatic-ring normal and the vector connecting the ring and CF_3_ centers ^18^. The pair of F201^5.62^ -CF_3_ from the tag sampled a predominantly perpendicular, shielding-favored geometry (mean θ = 27.30 ± 13.22°; 97.5% below the 54.7° magic angle; mean G = −1.619 ± 0.886 × 10^22^ cm^-3^). The corresponding F201^5.62^ contribution was negligible in the NECA-bound active-start trajectory (mean G = −0.001 × 10^22^ cm^-3^), in which H230^6.32^ contact was more orientationally heterogeneous. Its mean θ was 64.96 ± 14.25°, 75.9% of frames sampled the in-plane/deshielding side of the magic angle, and the ensemble-averaged G was +0.203 ± 0.433 × 10^22^ cm^-3^; H230^6.32^ contributed negligibly in the ZM-bound inactive-start trajectory (mean G = −0.014 × 10^22^ cm^-3^). Thus, in the inactive structure, the BTMB ensemble is influenced by a persistent F201^5.62^-associated shielding geometry, whereas in the active structure, the CF_3_ group of BTMB is closer to H230^6.32^.

A snapshot-level electronic-structure calculation provided an orthogonal qualitative check on the direction of this response (Fig. 5d). Expanded-region quantum-mechanical (QM) calculations were performed on representative frame-1 snapshots from BTMB- and BTFMA-labelled inactive and active models. For BTMB, isotropic shielding decreased from 242.072 ppm in the inactive snapshot to 240.293 ppm in the active snapshot (Δσ_active−inactive_ = −1.779 ppm), corresponding to an approximate downfield change of +1.779 ppm under Δδ ≈ −Δσ. BTFMA showed the opposite snapshot-level trend, with shielding increasing from 218.969 to 220.391 ppm (Δσ_active−inactive_ = +1.422 ppm), corresponding to an approximate upfield change of −1.422 ppm (Fig.5d).

To further assess whether these contacts were dictated by receptor conformation rather than influenced by ligand interactions, we removed ZM or NECA from the corresponding equilibrated structures and performed reference simulations. F201^5.62^ remained proximal to BTMB in the inactive-start control (4.84 ± 0.59 Å; mean G = −1.64 × 10^22^ cm^-3^), whereas H230^6.32^ remained proximal in the active-start control (4.89 ± 0.26 Å), suggesting that contact identity is largely determined by receptor conformation. Removal of NECA shifted the mean H230^6.32^ angle from 64.9° to 51.4° and the mean G value from +0.22 to −0.21 × 10^22^ cm^-3^, indicating that ligand binding can further tune the ring-current geometry. Overall, these state-dependent aromatic environments are consistent with the experimentally observed upfield enhancement with ZM and downfield enhancement with NECA, which lead to conformational shifts, respectively.

## Conclusion

We report a novel aryl-CF_3_ reporter, BTMB, with an increased solvent-dependent chemical-shift sensitivity and a structurally interpretable conformational-ensemble readout for the A_2A_R receptor. We fully characterized its susceptibility to microenvironmental change, structure-function perturbation when attached to the receptor, and the potential mechanism behind its superiority over the state-of-the-art probe, BTFMA, using MD simulations. This comprehensive investigation not only offers a novel probe to the community but also provides a strategy for continuing to explore next-generation ^19^F probes, as identifying ^19^F tags that best reflect the membrane protein’s conformational profile remains an ongoing task. Understanding why the new probe is better than the existing one, through MD simulations, is particularly important for guiding future probe discovery from a mechanistic perspective. Improving ^19^F-qNMR techniques to achieve high conformational resolution is essential for developing q-CAR-based drugs that specifically target disease-specific protein conformations^34–36^.

## Materials and Methods

### Comparative probe screening

Candidate fluorinated compounds were purchased from Sigma-Aldrich or Thermo Fisher Scientific and used as received without further purification. Their ^19^F chemical-shift sensitivity to changes in solvent polarity was evaluated using a methanol/water polarity series adapted from a previously established screening method^24^. Methanol and water were mixed at different volume ratios spanning relatively nonpolar to polar conditions, with MeOH/H2O ratios ranging from 4:1 to 1:4. Solvent polarity was expressed using the pyrene polarity scale (Py), with the corresponding Py values for the methanol/water mixtures taken from previously reported calibrations^37,38^.

^19^F NMR spectra were acquired at 20 °C on a Varian Inova spectrometer operating at a 600 MHz equipped with a dedicated ^19^F probe. The chemical shifts were referenced to NaF at −122.25 ppm. The chemical-shift change at each polarity was calculated relative to the least polar condition investigated, corresponding to MeOH/H2O = 4:1 (Py = 1.454), according to: Δδ(Py) = δ(Py) - δ(Py,ref), where δ(Py) is the observed ^19^F chemical shift at a given solvent polarity and δ(Py,ref) is the chemical shift measured in the least polar reference condition. The resulting polarity-dependent chemical-shift profiles were fitted to the sigmoidal function: Δδ = A / [1 + exp(−(Py − x₀)/w)], where A represents the magnitude of the chemical-shift response over the polarity range examined, x₀ is the midpoint of the transition, and w describes the width of the transition, with 1/w reflecting the steepness of the response. Chemical-shift sensitivity was compared primarily on the basis of the observed Δδ range and the fitted amplitude A.

### A_2A_R expression, purification, and fluorine labeling

Human A_2A_R was expressed using the previously established A_2A_R_316-V229C construct containing an N-terminal FLAG epitope and a C-terminal polyhistidine tag^2^. A single colony of *Pichia pastoris* SMD1163 expressing A_2A_R_316-V229C was inoculated into 4 mL YPD medium and cultured at 30 °C for 12 h, followed by expansion in 200 mL BMGY medium at 30 °C for 30 h. Cells were then transferred into 1 L BMMY induction medium and cultured at 20 °C. Methanol was replenished to 0.5% (v/v) every 12 h, and cells were harvested 60 h after initiation of methanol induction. Cells were collected by centrifugation at 4,000 × g for 20 min and washed once with 50 mM HEPES, pH 7.4, containing 10% (v/v) glycerol. Cell pellets were resuspended in breaking buffer containing 50 mM HEPES, pH 7.4, 100 mM NaCl, 2.5 mM EDTA, and 10% (v/v) glycerol at a buffer-to-cell ratio of 4:1 and disrupted by three passes through a microfluidizer at 20,000 psi. Unbroken cells and cellular debris were removed by centrifugation at 8,000 × g for 30 min, and membranes were collected from the supernatant by ultracentrifugation at 100,000 × g for 1 h. The isolated membranes were solubilized in 50 mM HEPES, pH 7.4, 100 mM NaCl, 1% MNG-3, and 0.2% CHS at 4 °C with gentle agitation. Talon resin was then added to the solubilized membrane fraction and incubated for at least 2 h at 4 °C. The A_2A_R-bound resin was washed with 50 mM HEPES, pH 7.4, 100 mM NaCl, 0.02% MNG-3, and 0.01% CHS. BTMB or BTFMA was first dissolved in 100% methanol to prepare a 100 mM stock solution. The probe was then added directly to resin-bound A_2A_R V229C to a final concentration of 100 μM and incubated at 4 °C with gentle agitation. After 12 h, a second aliquot probe was added, and labeling was continued for an additional 12 h at 4 °C with gentle agitation. Following labeling, excess probe was removed by extensive washing with the same detergent-containing buffer, and labeled A_2A_R was eluted with 50 mM HEPES, pH 7.4, 100 mM NaCl, 0.02% MNG-3, 0.01% CHS, and 250 mM imidazole. The centrifugation, membrane isolation, Talon capture, and on-resin labeling workflow follows the previously established A_2A_R purification procedure.

### Gα_s_ and Gβγ proteins’ preparations

Full-length human Gα_s_ was expressed as an N-terminal His₁₀–MBP fusion from a pBAD expression vector in *Escherichia coli* DH10B. We positioned an engineered flexible linker and a tobacco etch virus (TEV) protease cleavage site between MBP and Gα_s_. Bacterial cultures were grown in 2×YT medium at 37 °C to an OD₆₀₀ of approximately 0.6, cooled to 20 °C, and induced with 0.1% (w/v) L-arabinose. Protein expression was continued for 24 h at 18 °C. Cells were harvested, washed with 50 mM HEPES, pH 7.4, and 100 mM NaCl, and lysed in buffer containing 50 mM HEPES, pH 7.4, 100 mM NaCl, 0.02% MNG-3, 0.001% CHS, 1 mM MgCl₂, 100 μM GDP, and 20 mM imidazole. Following clarification by centrifugation, the supernatant was incubated with TALON cobalt-affinity resin at approximately 2 mL resin per liter of culture for 2 h at 4 °C. Bound Gαs fusion protein was eluted with buffer containing 300 mM imidazole. The eluate was dialyzed into cleavage buffer containing 50 mM HEPES, pH 7.4, 100 mM NaCl, 0.02% MNG-3, 0.001% CHS, 1 mM MgCl₂, 100 μM GDP, 0.5 mM EDTA, and 1 mM DTT, and the His₁₀–MBP fusion tag was removed by overnight TEV protease digestion at 4 °C using a TEV:Gαs ratio of approximately 1:50–1:100 (w/w). The cleavage mixture was subsequently passed over TALON resin, and untagged Gαs was recovered in the flow-through. Purified Gαs was buffer-exchanged into 50 mM HEPES, pH 7.4, 100 mM NaCl, 0.02% MNG-3, 0.001% CHS, 1 mM MgCl₂, and 100 μM GDP, concentrated, aliquoted, flash-frozen in liquid nitrogen, and stored at −80 °C. Gβ₁γ₂ was expressed in insect cells and purified from the membrane-associated fraction. Cell pellets from approximately 1 L of culture were washed with 150 mL of buffer containing 10 mM Tris-HCl, pH 7.5, 100 μM MgCl₂, and 5 mM β-mercaptoethanol. Membrane material was resuspended in 80 mL of solubilization buffer containing 20 mM HEPES, pH 7.4, 100 mM NaCl, 1% sodium cholate, 0.2% MNG-3, 5 mM MgCl₂, 5 mM β-mercaptoethanol, calf intestinal phosphatase, and protease inhibitors and extracted for 2 h at 4 °C. Following clarification by centrifugation, the soluble fraction was incubated with TALON resin for 2 h at 4 °C. The resin was washed first with solubilization buffer and subsequently with 20 mM HEPES, pH 7.4, 100 mM NaCl, 0.05% MNG-3, 1 mM MgCl₂, 5 mM β-mercaptoethanol, calf intestinal phosphatase, and protease inhibitors. Gβ₁γ₂ was eluted with the same buffer supplemented with 300 mM imidazole. The eluate was concentrated and dialyzed overnight at 4 °C against at least a 100-fold excess of washing buffer. Final purification was performed by anion-exchange chromatography using a low-salt buffer containing 20 mM HEPES, pH 7.4, 40 mM NaCl, 0.05% DDM, 1 mM MgCl₂, and 100 μM DTT, with elution against the corresponding buffer containing 1 M NaCl. Fractions containing purified Gβ₁γ₂ were pooled, concentrated, aliquoted, and stored at −80 °C.

### ^19^F NMR spectroscopy and spectral analysis

NMR samples contained 20–50 µM BTMB-labeled A_2A_R in 50 mM HEPES, pH 7.4, 100 mM NaCl, 0.02% MNG-3, and 0.01% CHS, with 10% D_2_O in a final volume of approximately 280– 300 µL. One-dimensional ^19^F NMR spectra were acquired at 20 °C on a Bruker NEO 600 MHz NMR spectrometer equipped with a QCI-F CryoProbe at the Integrated Molecular Structure Education and Research Center (IMSERC), Northwestern University. Typical acquisition parameters were a 16 µs 90° excitation pulse, a 200 ms acquisition time, a 15 kHz spectral width, a 1 s repetition time, and 15,000–50,000 transients. Chemical shifts were referenced to bendroflumethiazide at −59.05 ppm. Spectra were acquired for the ligand-free receptor and the receptor saturated with ZM241385, LUF5834, or NECA. For transducer-coupling experiments, NECA-bound receptor was incubated with the indicated heterotrimeric G-protein components before NMR acquisition. For G-protein-containing NMR samples, MgCl₂ and GDP were present at final concentrations of 1 mM and 100 μM, respectively. Partially overlapping ^19^F resonances were analyzed by multicomponent line-shape fitting in MestReNova 14.2 with minimized errors, using a common set of eight operational components (P1–P8) across the compared conditions. Fitted peak areas were normalized to the total fitted area of each spectrum and are reported as relative spectroscopic populations. The P1–P8 components were treated as operational spectral populations rather than one-to-one assignments to unique receptor structures.

Transverse relaxation times (T₂) were measured for representative resonances in the apo BTMB-labeled A_2A_R spectrum using a Carr–Purcell–Meiboom–Gill (CPMG) pulse sequence. Spectra were acquired at total transverse evolution times of 3.35 ms. Peak intensities were fitted to a single-exponential decay in MestReNova 14.2: I(t) = I₀ exp(−t/T₂), where I(t) is the resonance intensity at evolution time t, I₀ is the extrapolated intensity at t = 0, and T₂ is the apparent transverse relaxation time. Apparent homogeneous linewidths were calculated from T₂ according to: Δν₁/₂ = 1/(πT₂)

### G-protein activation assay

Receptor-dependent G-protein activation was measured with the GTPase-Glo assay (Promega)^39^. Purified A_2A_R and heterotrimeric Gα_s_βγ were mixed in a final volume of 10 µL containing 50 mM HEPES, pH 7.4, 100 mM NaCl, 0.001% CHS, and 0.05% MNG-3. Gα_s_βγ was maintained at a final concentration of 300 nM, and purified receptor was added at the indicated concentration. For comparisons of fluorine labeling, unlabeled control, BTMB-labeled, and BTFMA-labeled A_2A_R preparations were assayed at same receptor concentrations under otherwise identical conditions. Receptor–G-protein mixtures were preincubated for 30 min at room temperature. GTP hydrolysis was initiated by addition of 10 µL of 2× GTP-GAP solution containing 10 µM GTP, 1 mM DTT, and the cognate GTPase-activating protein (GAP), followed by incubation for 120 min at room temperature. Subsequently, 20 µL of reconstituted GTPase-Glo™ reagent containing 5 µM ADP was added to each reaction, and samples were incubated for an additional 30 min at room temperature with shaking. Luminescence was developed by addition of 40 µL detection reagent followed by a 10 min incubation at room temperature. Luminescence was measured using a BioTek FLx800 microplate reader at 528 ± 20 nm. All measurements were performed in three independent experiments. Data were analyzed using GraphPad Prism 10.

### Mass-spectrometric validation of probe labeling

BTMB- and BTFMA-labeled A_2A_R_V229C samples were analyzed by LC/TOF-MS to assess covalent probe modification of the V229C-containing peptide. Prior to denaturation, residual accessible thiols were alkylated with freshly prepared iodoacetamide (IAM) at a final concentration of 25 mM for 20–30 min at room temperature in the dark. Samples were subsequently denatured with 8 M urea and reduced with 5 mM tris(2-carboxyethyl)phosphine (TCEP) for 20–30 min at 37 °C. Before proteolysis, dilute the urea to ≤1 M with 50 mM ammonium bicarbonate. We added sequencing-grade trypsin at an enzyme-to-protein ratio of 1:50–1:100 (w/w) and digested at 37 °C for 8–16 h. Digestion was terminated by addition of formic acid to a final concentration of 0.5–1%, and peptides were desalted using C18 material prior to MS analysis. Peptides were analyzed on an Agilent 6230 LC/TOF-MS operated in positive-ion mode. We processed data using Agilent MassHunter Qualitative Analysis with the Find by Formula algorithm and a mass tolerance of ±10 ppm. Expected probe-modified peptide masses were calculated using monoisotopic mass shifts of +188.0449 Da for BTMB and +201.0401 Da for BTFMA on cysteine. Probe modification was assessed by comparison of the observed accurate masses and isotope patterns with the calculated masses of the corresponding unmodified, IAM-modified, and probe-modified V229C-containing peptides.

### BTMB labeling kinetics by Ellman’s assay

The time-dependent labeling of A_2A_R_V229C by BTMB was quantified using Ellman’s assay to determine the amount of accessible free thiol remaining during the labeling reaction. Labeling reactions were prepared in a total volume of 650 µL containing 10 µM A_2A_R_V229C. BTMB was initially added to a final concentration of 200 µM, and an additional 200 µM BTMB was added at 6, 12 h during the 24 h labeling period. Samples were gently mixed before collection, and 50 µL aliquots were withdrawn every 3 h from 0 to 24 h for Ellman’s assay. An unlabeled A_2A_R_V229C control was processed in parallel under otherwise identical conditions.

DTNB (Thermo Fisher Scientific) was prepared immediately before use. A 10 mM DTNB stock solution was prepared in 100 mM sodium phosphate buffer containing 1 mM EDTA at pH 8.0. A 5 mM DTNB working solution was prepared by mixing the 10 mM DTNB stock solution 1:1 with receptor purification buffer containing 50 mM HEPES, pH 7.4, 100 mM NaCl, 0.02% MNG-3, and 0.01% CHS. For each time-point measurement, 50 µL of receptor sample was transferred to a microplate well, followed by addition of 3.33 µL of 5 mM DTNB working solution. Samples were mixed immediately and incubated for 15 min at room temperature before absorbance was measured at 412 nm. A reagent blank containing 50 µL receptor purification buffer and 3.33 µL of 5 mM DTNB working solution was measured in parallel and subtracted from the corresponding sample absorbance. Three independent labeling experiments were performed. Kinetic fitting and statistical analyses were performed using GraphPad Prism 10.

### Structural modeling and covalent construction of BTFMA- and BTMB-labeled A_2A_R

Inactive- and active-like receptor models were used from PDB 3EML and 6GDG, respectively^32,33^. Missing loops were rebuilt with MODELLER through UCSF ChimeraX^40,41^, and candidate models were inspected for steric clashes before introduction of V229C. BTMB was attached to C229 using the CHARMM-GUI covalent-ligand workflow^42^. Multiple poses were generated, and a chemically reasonable representative from the major cluster was selected for system preparation. Covalent attachment was represented by replacing the C– Br bond with a bond between the C229 sulfur and the benzylic carbon of BTFMA and BTMB. The resulting receptor–BTMB or BTFMA complexes were transferred to CHARMM-GUI for micelle-system preparation^43^.

### Molecular dynamics simulations

All-atom simulations were performed with NAMD 3.0.3 using the CHARMM36m framework^44,45^. ZM-bound inactive-like and NECA-bound active BTMB-A_2A_R systems were prepared under matched conditions in detergent micelles containing 100 BLMNG and 20 CHSD molecules, solvated with TIP3P water and approximately 0.10 M NaCl. Simulations were conducted at 293.15 K and 1 atm under periodic boundary conditions. Long-range electrostatics were treated using particle-mesh Ewald, and van der Waals interactions were switched from 10 to 12 Å. Each system underwent energy minimization and staged equilibration with progressive release of positional restraints on receptor, BTMB and orthosteric ligand. Production trajectories were propagated for 100 ns with a 2 fs timestep and constraints on hydrogen-bonded bonds. Coordinates were saved every 50 ps, 2,000 frames per 100-ns trajectory. Short 10-ns ligand-free controls were initiated by removing ZM or NECA from the corresponding equilibrated bound-state system and were interpreted only as local relaxation controls.

### Aromatic contact and ring-current geometry analysis

Aromatic contacts were analyzed with MDAnalysis^46^. For each frame, the centre of mass of the BTMB CF_3_ group and each protein aromatic ring (Phe, Tyr, Trp and His) was calculated using periodic minimum-image distances. Aromatic-ring planes were defined from ring heavy atoms, and θ was the acute angle between the ring normal and the vector connecting ring and CF_3_ centres. We excluded the aromatic ring of BTMB itself. The geometric factor G = (1 - 3cos^2^θ)/r^3^ was calculated using the CF_3_-to-ring centre distance r ^18^. The 54.7° magic angle separates negative and positive values of the geometric term. We interpreted negative G as a shielding/upfield tendency and positive G as a deshielding/downfield tendency. We summarized distributions over all saved frames; we did not convert G directly to ppm.

### Quantum-mechanical shielding calculations

Expanded-region ^19^F shielding calculations were performed on representative frame-1 snapshots from the BTMB- and BTFMA-labelled ZM- and NECA-bound models with ORCA 6.1.1 using the PBE0 functional and pcSseg-2 basis set with the GIAO formalism^47–50^. The QM region comprised the fluorinated reporter (BTMB or BTFMA) and the C229 side chain, truncated at Cβ and capped with hydrogen. Electrostatic embedding was represented by CHARMM point charges from complete residues and molecules within 12 Å of the CF_3_ fluorine atoms. The isotropic shieldings of the three fluorine nuclei were averaged within each snapshot. State-dependent chemical-shift changes were interpreted using Δδ ≈ −Δσ; absolute shielding values were not compared across reporter chemistries.

## Acknowledgements

This work is funded by the National Institutes of General Medical Sciences (1R01GM149659, L. Y.) and the National Institute of Environmental Health Sciences (1R21ES035378, L. Y.) and start-up funding from the University of South Florida. We also thank Wenjie Zhao for helping conduct some ^19^F-NMR experiments, and Dr. Heng Liu in the Department of Chemistry at the University of South Florida for his assistance with LC-MS data collection.

## Author Contributions

L. Y. conceptualized and designed the study. X.W. and Y.C. prepared and purified receptors and Gα protein constructs. W.Z. performed sample preparation and conducted ^19^F-NMR of free compounds. W. S. prepared Gβγ proteins. X. W. also performed NMR spectral acquisition, data processing, and spectral analyses in addition to biochemical assessments. Y. C also conducted the biochemical assessments and NMR data acquisition and data analyses. T. M. assisted in receptor preparation. Y. Z. assisted with setting up and optimizing the NMR pulse sequence parameters. L. Y. supervised the whole project. X. W. and L. Y. wrote the manuscript, and all authors contributed to the revisions.

